# High total water loss driven by low-fat diet in desert-adapted mice

**DOI:** 10.1101/2022.04.15.488461

**Authors:** Danielle M. Blumstein, Jocelyn P. Colella, Ernst Linder, Matthew D. MacManes

## Abstract

Availability of food resources is an important driver of survival. Animals must either relocate or adapt to persist in environments where critical resource abundance is changing. An optimal diet balances energy gain, water regulation, and nutrition. We used flow-through respirometry to characterize metabolic phenotypes of the desert-adapted cactus mouse (*Peromyscus eremicus*) under diurnally variable environmental conditions that mimic the environment of the Sonoran Desert. We treated mice with two different energetically equivalent diets, a standard diet and a low-fat diet, and measured energy expenditure, water loss rate, respiratory quotient, weight, and electrolyte levels. Mice fed the low-fat diet lost significantly more water than those on the standard diet. Our results suggest that cactus mice may have limited capacity to tolerate water deprivation if optimal foods become less abundant. Given that climate change is predicted to modify the distribution of food items, understanding these links may have important implications for long-term population viability for desert and non-desert adapted animals alike.

## Introduction

One of the most critical challenges faced by desert mammals is the regulation of water. Water balance is maintained through a cascade of hormones that elicit a response designed to increase water retention and decrease water use in the excretion of nitrogenous wastes. Dehydration, in contrast, is an outcome of inadequate water balance. This results in a decrease in blood volume and an increase in osmolality, the latter, an effect driven largely by the rise in serum sodium (Thornton 2010; Leib et al. 2016) and eventually leading to death if water homeostasis is not achieved. While hormonal regulation of water balance is paramount to survival, behavioral, physiological, and genomic responses are also critical.

Dietary intake of food is one important way in which animals manage water. There are three main macromolecules that animals ingest as food items; carbohydrates, fats, and proteins, all of which vary in their amount of energy and water, as well as in their cost to metabolize. Energy potential varies greatly, with carbohydrates and proteins yielding 16.74 J/g, while fats yield more than double that (37.66 J/g, Sánchez-Peña et al. 2017). Metabolic water, the water produced through endogenous catabolism of carbohydrates, fats, or proteins (Schmidt-Nielsen and Adolph 1964; Frank 1988; Orr et al. 2015 production also varies dramatically. Oxidation of carbohydrates yields 0.60 g of metabolic water per gram and fats yield 1.07 g per gram due to greater oxygen requirements of lipid metabolism, while proteins yield the least metabolic water (0.41 g) of any macronutrient and even require water loss for excretion of nitrogenous waste (Mellanby 1942; Davidson and Passmore 1963; Schmidt-Nielsen and Adolph 1964).

In environments like deserts, where extrinsic water is limited, both preformed (dietary) water and endogenous water production are key to survival. Dietary intake represents the raw material used to produce metabolic water. Food catabolism results in water loss through respiration, urination, and fecal production (Schmidt-Nielsen 1975). Specific dietary composition is often unknown and highly variable, particularly for wild animals, energy potential (Pyke et al. 1977; Pyke 1984), nutrition, and capacity for water production (Schmidt-Nielsen and Adolph 1964) all depend on diet composition. In deserts, vegetation, seeds, and insects comprise the bulk of rodent diets (Reichman 1975) with each varying in fat, protein, and carbohydrate composition and seasonal availability (Wolf and del Rio 2003; Orr et al. 2015). In response to natural variation in resource availability, animals have the opportunity to make dietary decisions that may have a substantial effect on their internal water economy.

Mice of the genus *Peromyscus* have the widest distribution of any North American mammal and are considered model organisms in evolutionary biology due to their unparalleled habitat diversity (Bedford and Hoekstra 2015), extensive genomic resources (*e.g.,* Tigano et al. 2020; Colella et al. 2021a), abundance of historical and contemporary collections in natural history museums (Pergams and Lawler 2009; Pardi et al. 2020), and ability to live and breed under laboratory conditions (Crossland et al. 2014). The desert specialist cactus mouse, *Peromyscus eremicus*, is endemic to the southwestern United States and exhibits behavioral (Murie 1961; Veal and Caire 1979), physiological (Macmillen 1965; Kordonowy et al. 2017; Colella et al. 2021b), and molecular (MacManes 2017; Tigano et al. 2020) adaptations to desert environments, making it an interesting natural experimental model to examine mechanisms of adaptation to warmer, drier environments. *Peromyscus eremicus* are omnivorous and opportunistic in their diet, utilizing seeds, arthropods, and green vegetation seasonally (Bradley and Mauer 1973; Meserve 1976) and they shift to consuming cactus seeds and/or fruit pulp during summer months (Orr et al. 2015).

We examine the effect of dietary fat content on the metabolic physiology of the desert-adapted cactus mouse to understand the physiological consequences of food availability and dietary choices. To accomplish this, we extend previously characterized circadian metabolic patterns for males and females (Colella et al. 2021b) to estimate rates of metabolism and water loss in animals fed an experimental diet low in fat, but unchanged in terms of energy composition and level of macronutrients, to those on a standard laboratory diet. This experimental framework allows us to test the relationships between dietary fats, energy expenditure, and water balance, which have direct connections to fitness. Given that fat catabolism yields more water and energy compared to carbohydrates or proteins, we hypothesized that mice fed a low-fat diet would perform more poorly than those on the standard diet.

## Methods

### Animal Care and Experimental Model

We worked with a captive colony of cactus mice maintained at the University of New Hampshire. Captive animal care procedures followed guidelines established by the American Society of Mammalogists (Sikes et al. 2016), American Veterinary Medical Association (Leary et al. 2013), and were approved by the University of New Hampshire Institutional Animal Care and Use Committee (IACUC, #103092).

Sexually mature, non-reproductive healthy adult mice (n = 28 males, n = 28 females) between three and nine months of age, bred from wild-derived lines at the *Peromyscus* Genetic Stock Center at the University of South Carolina (Columbia, South Carolina, USA) were used in this study. Mice were randomly assigned to one of two feeding groups: a standard diet group (SD - LabDiet® 5015*, 26.10% fat, 19.752% protein, 54.15% carbohydrates, energy 15.02 kJ/g, food quotient [FQ, the theoretical respiratory quotient produced by the diet based on macronutrient composition, Westerterp, 1993] 0.89) or a low-fat diet group (LFD - Modified LabDiet® 5015 [5G0Z], 6.6% fat, 22.8% protein, 70.6% carbohydrates, energy 14.31 kJ/g, FQ 0.92). All food was stored in the desert chamber to match the environmental conditions described below to control for the water content in the diets. Animals were housed individually in 9.5 L animal chambers inside a larger, environmentally controlled room built to simulate the temperature, humidity, and photoperiodic conditions of the Sonoran Desert (Kordonowy et al. 2017; Colella et al. 2021b). Each animal chamber contained dried cellulose-based bedding, and animals were provided food and water *ad libitum*. Mice were acclimated to the LFD for one month and to experimental cages for 24 hours prior to the beginning of metabolic measurements. Animals were weighed to the nearest tenth of a gram on a digital scale before being housed in an experimental chamber for four days during metabolic data collection. Animals were weighed again at the end of the experiment.

Environmental temperatures and relative humidity (RH) followed a natural, diurnal pattern. Temperature and RH were fixed at 32°C and 10% RH during the light phase for 11 hours (06:00 to 20:00) and dropped to 24°C and increased to 25% RH during a one-hour transition starting at 20:00. Temperature and RH then remained constant during the dark phase for seven hours, before the temperature increased again and the RH fell to light phase temperature during a three-hour transition that started at 06:00. Photoperiod was regular with 16 hours of light and 8 hours of dark. See Figure 1A for a visual representation of environmental conditions.

**Figure 1.**
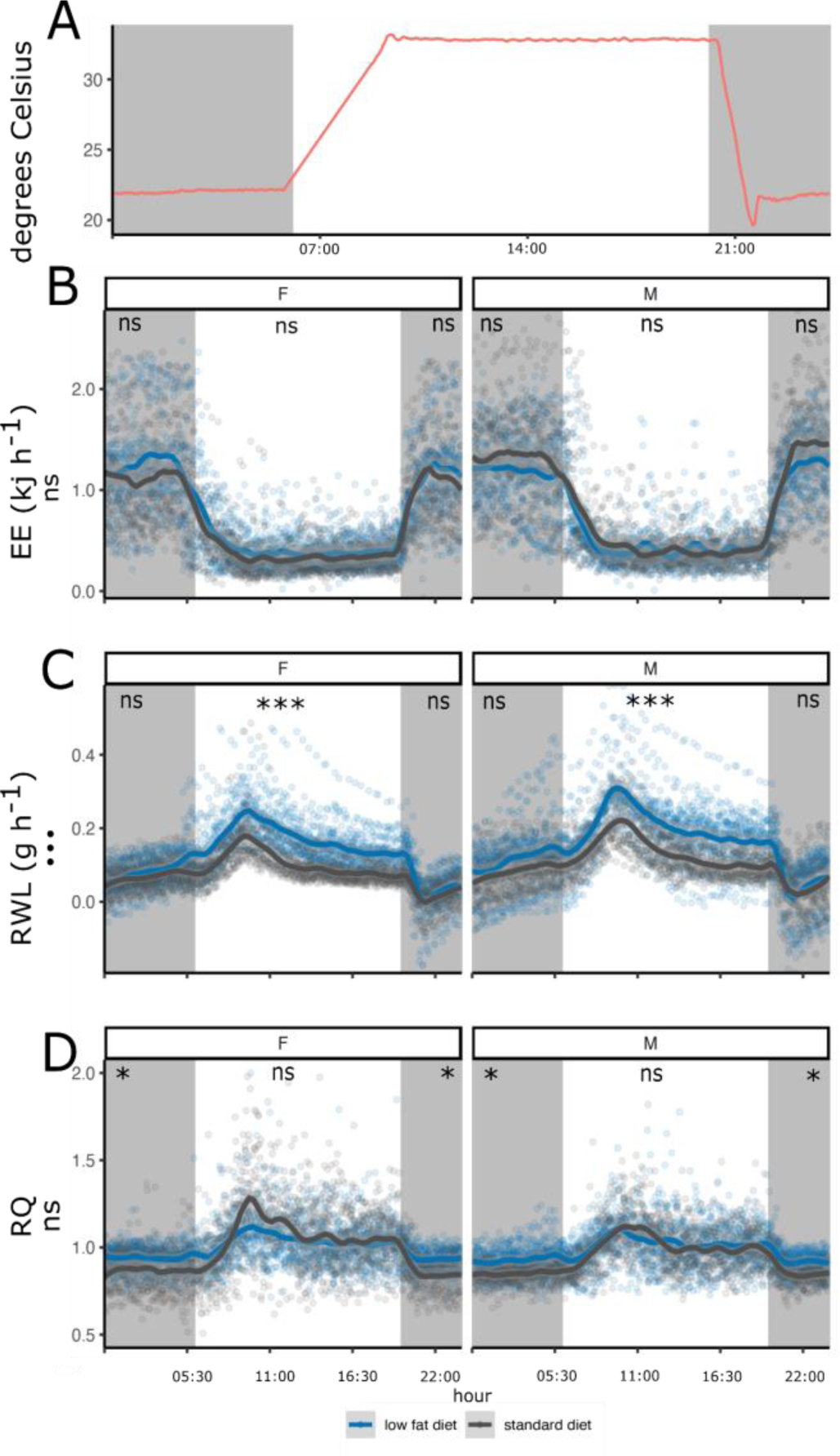
72 hours of environmental (A) and respirometry (B-D) data split by sex for 28 adult males and 28 adult females plotted over a 24-hour window to display circadian patterns for each group. Organismal responses to standard diet are in gray and low-fat diet in blue. Vertical blocks shaded in gray indicate the dark phase when animals are active, and unshaded blocks indicate light phase when animals are inactive. **A)** Room temperature (degrees Celsius) is indicated by a solid red line. **B)** 24-hour measurements of energy expenditure (EE kJ hr^-1^), **C)** water loss rate (WLR, H_2_O g hr^-1^), and **D)** respiratory quotient (RQ), for females (F, left) and males (M, right). Statistical significance between the fixed effects, diet and sex, for the 24hr GAMMs are denoted with one or more · under the y-axis label, ‘ns’ indicates non-significance. Statistical significance between the fixed effects, diet and sex, from GAMMs on the subsetted day/night data are denoted with one or more * on the top axis of the plots. The two symbols (*/·) signify the same significance values (* p <= 0.05, **: p <= 0.01, ***: p <= 0.001, ****: p <= 0.0001).

### Metabolic phenotyping

Metabolic phenotyping data were collected starting at 12:00 using a previously described (Colella et al. 2021b) commercially available Field Metabolic System purchased from Sable Systems International (SSI). Physiological data were collected following the methods outlined in Whitfield et al. (2015) and Colella et al. (2021a). Oxygen consumption (VO_2_ ml min^-1^), carbon dioxide production (VCO_2_ ml min^-1^), and water vapor pressure (WVP, kPa) lost via urine, feces, cutaneous, and respiration were continually measured during 72-hour trials. The experiment was repeated twice for each sex under each diet (SD and LFD), resulting in three-day experiments performed sequentially for eight batches of seven mice (and one empty, baseline chamber) per three-day experiment. Batches alternated between females and males for a total of 28 adult males and 28 adult females.

Airstreams were pulled from flow-through chambers at a constant rate of 1600 ml min^−1^ (96 L h^−1^) by an SSI SS-4 sub-sampler pump (one for each chamber), multiplexed through an SSI MUXSCAN, sub-sampled at 250 ml min^−1^ into a Field Metabolic System (FMS) where WVP, CO_2_, and O_2_ were analyzed with no scrubbing. The FMS was zeroed, and sensors were spanned between each 72-hour experiment following the SSI Instrument Settings and Calibration manual using dry gas of known CO_2_ and O_2_ concentrations.

To generate accurate individual measurements, chambers were measured in a pseudorandom order for 120s each, approximately three times every hour. Using methods described in Lighton (2018), we used the most stable 50% of each 120s period to produce a single averaged value for the rate of O_2_ consumption (ml min^-1^), rate of CO_2_ production (ml min^-1^), and water loss rate (WLR) (mg h^-1^) per measurement window. Lag time between the analyzers was corrected so that measurements represent the same time periods (Lighton 2018). This procedure results in a highly repeatable set of measurements. VO_2_ and VCO_2_ were calculated using equations 10.5 and 10.6, respectively, from Lighton (2018). Respiratory quotient (RQ) was calculated as the ratio of VCO_2_ to VO_2_ (Lighton 2018). Energy expenditure (EE), kJ hr^-1^, was calculated as in Lighton (2018, eq. 9.15). Response variable measurements that were more than three standard deviations away from the mean, which often represented times when animal care staff entered the room, were considered outliers, and therefore were removed from downstream analysis.

### Electrolytes

At 12:00 Eastern Standard Time (EST), immediately following the conclusion of the experiment, animals were weighed and euthanized with an overdose of inhaled isoflurane and decapitation. 120 µL of trunk blood was collected within one minute of death for serum electrolyte measurement using an Abaxis i-STAT® Alinity machine and CHEM8+ cartridges (Abbott Park, IL, USA, Abbott Point of Care Inc.), and calibrated automatically with each cartridge run. We used CHEM8+ cartridges to measure the concentration of sodium (Na), potassium (K), creatinine (Cr), blood urea nitrogen (BUN), hematocrit (HCT), glucose (Glu), and ionized calcium (iCa), which are expected to vary in response to hydration status and renal function. Finally, using sodium, glucose, and BUN, we calculated osmolality using the formula in Rasouli (2016). After assessing for normality, we used a student’s two-tailed t-test in R (stats::t.test) to test for significant (p < 0.05) differences in weight and electrolytes within and between the treatments and sexes for each experimental group.

### Statistical Analysis

All statistical analyses were conducted in R v 4.0.3 (R Core Team 2020) unless otherwise specified. To perform analyses on the shape of the temporally variable curves, we modeled the effect of time in days, diurnal time, and EE, RQ and/or WLR on EE, RQ, and WLR using Generalized Additive Mixed Models (GAMMs) using the gamm function in the mgcv R package (Lin and Zhang 1999; Wood 2017). To avoid violating the assumption of independence, we used the experimental batches and mice within each experimental batch as random effects. The R function gamm then simultaneously estimates the variances of these random effects as well as the treatment effects, thus implicitly adjusting treatment and covariate effects for the effect of correlated, nested groups of measurements. As a result, treatments do not explain differences between individual mice, but rather the average difference between groups of mice, because we were interested in the diet treatment and the effect of sex, as well as in the response over the full 72-hour period and the diurnal cycle. Our models included two fixed effects, diet and sex, and four nonlinear smoothing regression terms: time in days as a nonlinear trend, diurnal cycle as a circular spline function, and two of the three respirometry response variables (EE, RQ, and/or WLR). Using the effective degrees of freedom (edf), we were able to determine the degree of non-linearity of each curve (Wood 2017). To test for physiological differences between mice fed each diet during the light and dark phases, we subsetted data collected during the light phase and dark phase for each response variable and ran GAMMs as described above.

To test for differences in the total volume of water lost for each animal over the course of the experiment, rather than the shape of the curve, we estimated the amount of water lost per individual for every hour of the experiment and summed those values. These values were then used as the response variable in an ANCOVA (car::anova, Fox and Weisberg 2019). The model consisted of two independent variables, sex and diet, and the interaction between them. Weight at the start of the experiment was used as the covariate. Pairwise comparisons were made using the Tukey separation of adjusted means test (emmeans:: emmeans and emmeans::contrast, Lenth 2021).

## Results

Data from 56 sexually mature (but non-reproductive) male and female mice were collected as described above. No health issues were detected by veterinary staff during the duration of the experiment, nor were there significant differences in weight between males and females (mean of 22.20 g [range: 15.40 - 29.10 g] versus 21.2 g [range: 16.91 - 29.34 g], respectively), or between diet groups.

### Metabolic Phenotyping

In total, we recorded 8,125 120s intervals: 4,091 female, 4,034 male (Supplemental Table 1). We measured metabolic variables for 4,105 120s intervals under the LFD: 2,053 female and 2,052 male measurements. For the SD treatment, we recorded 4,020 total observations (2,038 female, 1,982 male). After removing outliers, we retained 3,945 female measurements (1,977 SD and 1,968 LFD) and 3,963 male measurements (1,945 SD, 2,018 LFD). *Peromyscus eremicus* show physiological patterns that are in phase with photoperiod and room temperature for both diets (Figure 1), as described in (Fox and Weisberg 2019; Colella et al. 2021b). Raw data and processed machine-readable csv files are available on Zenodo: https://zenodo.org/record/6422231#.YlXSD9PML0o.

### Energy Expenditure

Consistent with previous work (Colella et al. 2021b), males and females show diurnal patterning of EE on both diets, with the highest EE occurring during the dark phase when environmental temperature is lowest and animals are active. The lowest EE occurred during the light phase, when environmental temperature was greatest and animals were inactive (Figure 1B). Using a GAMM, we modeled EE with diet and sex as fixed effects, as well as other predictors, such as day, time, WLR, and RQ, and found that neither sex nor diet influenced EE (diet p = 0.56, sex p = 0.18, Supplemental Table 2). When analyzing light and dark phases separately, there were no differences in EE between males and females nor between the two diets (light phase: diet p = 0.550, sex p= 0.293; dark phase: diet p = 0.571, sex p = 0.182, Supplemental Table 3 and 4). There was a weak nonlinear relationship between EE and light phase (Supplemental Figure 2A, Supplemental Table 2, edf = 1.98, p < 2e^-16^). Additionally, the relationship between time and EE was statistically significant but nonlinear (Supplemental Figure 2F, Supplemental Table 2, edf = 7.98, p < 2e^-16^).

### Water Loss Rate

Both treatment groups, SD and LFD, and both sexes exhibited diurnal patterning of WLR and had the highest WLR during the light phase when environmental temperature was highest, even though animals were inactive then (Figure 1C, Supplemental Figure 1F, Supplemental Table 2, edf = 7.95, p < 2e^-16^). Peak loss occurred between 09:00 and 11:00. By fitting a 24-hour time-continuous GAMM (Supplemental Figure 1E-H), we showed that both diet and sex had a significant effect on the WLR (diet p = 2.13E^-05^, sex p = 0.002, Supplemental Table 2), with both males and females fed a LFD losing more water than those fed the SD. When modeling water loss during the light phase only, we showed that both diet and sex had a significant effect on WLR (diet p = 4.80e^-15^, sex p = 1.69e^-11^, Supplemental Table 3). When modeling water loss only during the dark phase, when animals are active, WLR was significantly different between males and females (p = 0.015, supplemental table 4), but WLR was not different within sex between different diets (p = 0.797, Supplemental Table 4). The relationship between WLR and experimental day was weakly non-linear, but significant (Supplemental Figure 1E, Supplemental Table 2, edf = 1.99, p < 2e^-16^).

Both diet and sex significantly affected the amount of water lost over the course of the experiment when controlling for individual body weight (p = 5.22e^-2^ and p = 0.017, respectively, Supplemental Table 5), but the interaction term was not significant (p= 0.633, Supplemental Table 5). Further investigation into the sex:diet interaction showed significant differences in both the female (p=0.043, Supplemental Table 5) and male (p=0.005, Supplemental Table 5) diet comparison.

### Respiratory Quotient

RQ showed circadian rhythms for both diets and sexes with the highest RQ occurring during the light phase and the lowest during the dark phase (Colella et al. 2021b; Figure 1D). By fitting a 24-hour GAMM we found no significant difference in RQ between the sexes or diets (sex p = 0.83, diet p = 0.52, Supplemental Table 2) but we did find that day and time were highly significant (p = 1.98e^-07^, p < 2.2e^-16^ respectively). Time of day had a complex relationship with RQ (Supplemental Figure 1, edf = 7.69, Supplemental Table 2) and experimental day had a weak non-linear relationship with RQ (Supplemental Figure 1D, edf = 1.59, Supplemental Table 2).

During the light phase, RQ rose above the FQ, with females fed the SD having the highest RQ at the start of the light phase and then stabilizing at an RQ > 1 around 14:00. RQ did not differ by sex nor diet (sex p = 0.740, diet p = 0.543, Supplemental Table 3). When modeling RQ during the dark phase, values were comparable to the FQ of the respective diet for both sexes (FQ LFD: 0.92, FQ SD: 0.89) and RQ was significantly different between the diets (p = 0.017, Supplemental Table 4) but not between males and females (p = 0.739, Supplemental Table 4).

### Electrolytes

Several serum electrolyte values were significantly different between diets for males and females (sodium p = 0.22 and p = 0.01 respectively; potassium p = 0.02 and p = 0.01; BUN p = 0.03 and 0.71; hematocrit p = 0.04 and 0.02; ionized calcium p = 0.001 and 0.001; osmolality p = 0.01 and 0.05) (Figure 2, Table1). When comparing males and females on the same diet, sodium (p = 0.02) and Osmolality (p = 0.03) differed significantly for mice on the SD and no electrolyte measurements were differed between males and females on the LFD.

**Figure 2.**
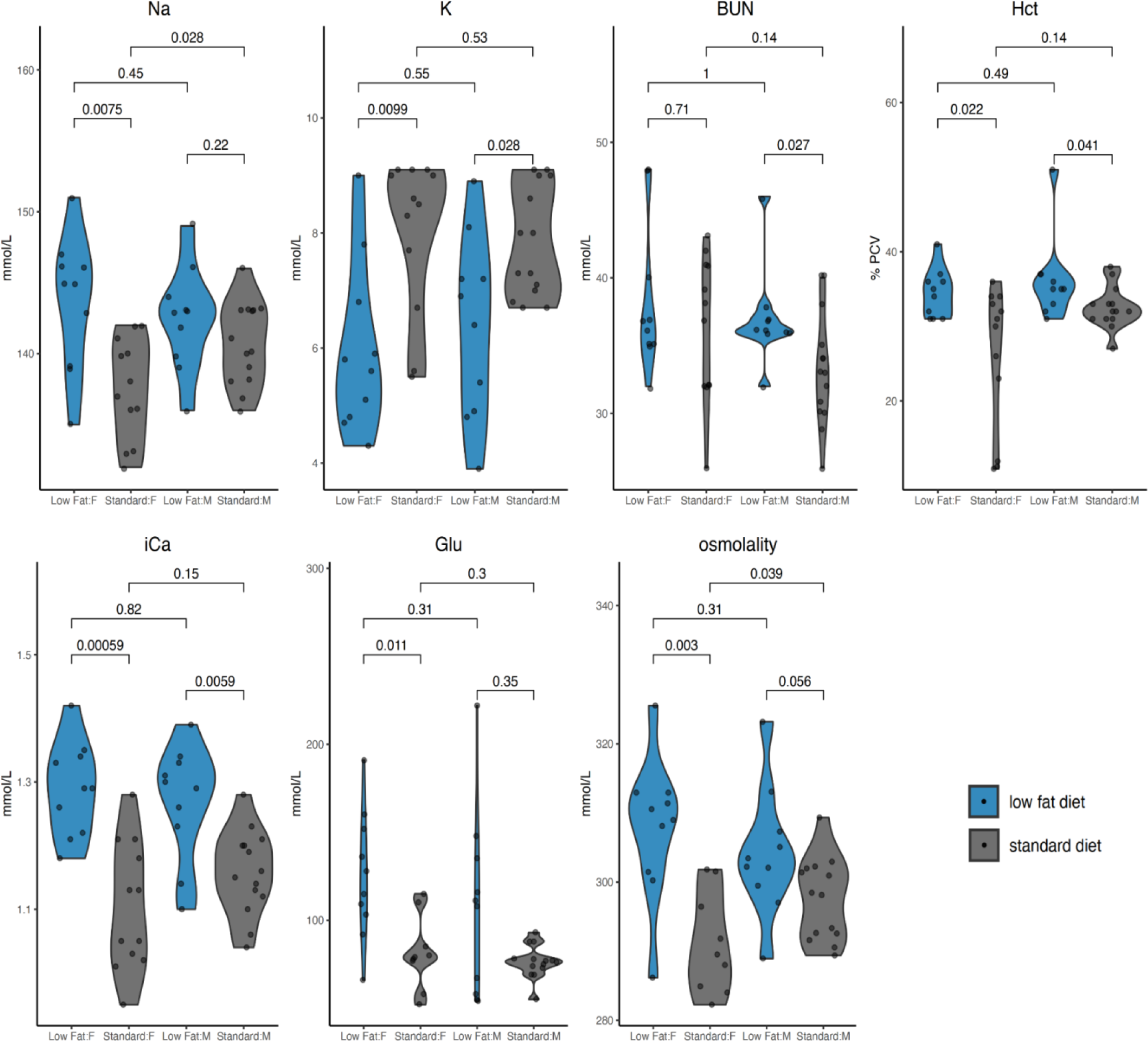
Violin plots showing the distribution of serum electrolyte measurements: Na = sodium (mmol/L), K = potassium (mmol/L), BUN = blood urea nitrogen (mmol/L), Hct = hematocrit (% PCV), iCa = ionized calcium (mmol/L), Glu = glucose (mmol/L), and osmolality (mmol/L) for female (F, left) and male (M, right) *Peromyscus eremicus* fed standard (grey) and low-fat (blue) diets. Observations (28 adult females and 28 adult males, n=14 of each treatment, total n=56) are represented by black dots. P-values from pairwise t-tests are reported above the brackets.

## Discussion

We explored the effect of dietary fat intake on water balance in a desert adapted rodent and found that mice fed a diet lower in fat lost more water than mice fed a diet higher in fat. In environments where water scarcity is extreme, even small differences in water balance could have substantial impacts on cognitive function and physical performance, and therefore survival and reproductive success (Boogert et al. 2018). Our results help clarify the physiological consequences of food availability and dietary choices.

### Energy Expenditure

Not all diets provide animals with the same amount of energy per unit mass (Withers 1982). Different macronutrients have different energy potentials (Sánchez-Peña et al. 2017). Diet composition influences EE (Acheson 1990; Westerterp 2004; Secor 2009; Beale et al. 2018), as do other intrinsic (*i.e.,* body mass, temperature, evolutionary history, reproductive stage, activity, or energy intake) and extrinsic (*i.e.,* habitat [including environmental temperature], climate, or social environment; Speakman 1997) variables. In wild populations, dietary switches tend to coincide with seasonal changes that affect resource quantity and/or quality (Noble et al. 2019). Previous studies have recorded cactus mice shifting diet seasonally, consuming arthropods during the winter (Hope and Parmenter 2007), and transitioning to the consumption of cactus seeds and/or fruits during the summer (Hope and Parmenter 2007; Orr et al. 2015). Many plants native to the Sonoran Desert have a fat content comparable to that of the LFD used in this study (5.49 - 9.88% for leaf tissues and branches, McArthur 1994; 0.90-3.60% for roots and seeds, Castle et al. 2020).

In the current study, both diets were designed to have equivalent amounts of energy per gram of weight. We designed the experiment to control for several intrinsic and extrinsic factors that represent sources of variation in EE, but not in individual activity. We show that experimental manipulation of diet does not result in differences in EE when food is provided *ad libitum*. EE and activity are tightly coupled (Garland et al. 2011; Kaiyala et al. 2012). Therefore we expect, at most, limited differences in activity given limited differences in EE. For both diets, animals are likely to eat enough to satisfy energy needs, a hypothesis further supported by the fact that there was no net change in weight over the course of the experiment. Consistent with other studies, we recorded higher EE during the dark phase of the 24-hour cycle when animals were active (Colella et al. 2021b), further supporting that metabolic rate is complex and may be modulated by the environment, activity, or other extrinsic/intrinsic factors.

### Water Loss Rate

#### Thermoregulation and water loss

Evaporative water loss is one way for organisms to cool themselves (*e.g.,* via latent heat of vaporization) and results in a decrease in body temperature and loss of body mass (Porter and Gates 1969). While body temperature was not measured as a part of this study, numerous studies have shown that desert animals reduce respiratory water loss as an adaptive physiological mechanism for maintaining water balance in desert environments (Schmidt-Nielsen and Schmidt-Nielsen 1951; Hart 1971; MacMillen 1972). Indeed, a fine balance must exist between using water for thermoregulation, the use of water for other critical metabolic processes, and dehydration. Exaggerated use of water for thermoregulation may be harmful when water is limited. Instead, other less water-intensive mechanisms of heat loss such as conduction, and behavioral changes such as estivation and nocturnality may be used to maintain a homeostatic body temperature but are not quantified in this study.

Despite the consequences of limited water supply, desert *Peromyscus* have significant capacity for evaporative cooling for thermoregulation (Ramirez et al. 2022). In our study, we found that animals fed the LFD had greater water loss compared to animals fed the SD (Figure 1C). While that loss may be related to differences in thermoregulatory performance, it may further compromise water balance. Are these patterns suggestive of differences in physiological performance? Is the consumption of a diet lower in fat related to a reduction in physiological performance and, consequently, fitness? While the current experiment cannot definitively answer these questions, there is some evidence (*e.g.,* electrolytes, water loss) to suggest that physiological performance is impaired in animals consuming the LFD and that this performance, particularly when water is scarce, may have fitness consequences.

#### Potential mechanisms for increased water loss on LFD

Across all comparisons, there is a strong and statistically significant relationship between the water loss rate and dietary fat content, with individuals fed the LFD losing more water. While the potential mechanisms underlying these findings will require future experiments, several can be ruled out. First, the maintenance of water balance is a process regulated by a cascade of hormones including arginine vasopressin and the renin–angiotensin–aldosterone system pathway (Aisenbrey et al. 1981; Greenleaf 1992). While some hormones are lipid-based (*e.g.,* aldosterone) and their production and secretion are linked to dietary composition (*i.e*., sex hormones and severe low body fat), the degree to which dietary fats are reduced for mice fed the LFD are not to the level at which hormonal imbalances should be seen. Second, while there is a known relationship between ketogenic diets and dehydration (Wheless 2001; Freeman et al. 2006), these diets are associated with enhanced diuresis. We observed the opposite with animals fed the SD; they lost less water than animals fed the LFD. Third, diets higher in protein require more water for the removal of nitrogenous wastes (Calder and Braun 1983), but higher protein does not account for the observations herein. To approximate equivalent energy availability there is a small difference in the amount of dietary protein in each dietary treatment (18.9% in SD versus 19.5% in LFD), however, that small difference is unlikely to result in the observed differences in water loss and hydration status.

While food and water intake were not measured directly, a potential mechanism for the observed differences could be that animals fed the LFD simply eat more than animals fed a diet higher in fat. While the diets are closely matched in terms of available energy per unit mass, the LFD is slightly less energy dense. Additionally, it’s possible that there are substantial differences in the non-nutritive fraction of the diet that could be further explored. While the current study does not disentangle this potential aspect, future work could. These differences suggest that animals fed the LFD may be ingesting more food mass to maintain nutritional homeostasis and with food intake increased there may also be increased water intake (Bachmanov et al. 2002). While we have no evidence for this, an increased rate of consumption would potentially result in an increased rate of urine production and the production of water-containing fecal material and urine. Given that animals fed the LFD seem to be more dehydrated than animals fed the SD (increased Na, osmolality, Hct, Figure 2), this hypothesis requires that animals fed the LFD consume more water, but not enough to satisfy their physiological needs. Given that the animals have free access to water, an unmet need is irreconcilable without further explanation.

Another intriguing hypothesis linked to the concept of unmet hydrational needs has to do with thirst pathways. Thirst is a complex phenotype under tight neural control. Osmo- and baroreceptors located in the vasculature are responsible for monitoring water balance and, when triggered, set into motion a hormonal cascade that simultaneously enhance renal water retention and stimulate thirst (Thornton 2010; Leib et al. 2016). While the former aspects of the response are crucial to many physiological processes, the latter response, thirst, could be responsible for the observations described above. While purely a hypothesis, there are several examples of species losing aspects of their sensory systems via natural selection, loss of sight (Gore et al. 2018), decreases in certain taste sensitivities or loss of taste entirely (Jiang et al. 2012), and loss of specific olfactory functions (Kishida et al. 2015). For animals in dry deserts, where water is scarce, could well-developed pathways leading to thirst be similarly changed? While requiring careful consideration, perhaps using a comparative genomics approach, this hypothesis could explain why animals with free access to water remain significantly dehydrated.

### Respiratory Quotient

RQ, the ratio of CO_2_ produced to O_2_ consumed, can inform rates of differential fuel utilization. RQ values typically fall between 0.7 and 1.0, with the catabolism of fats yielding 0.7, carbohydrates 1.0, and proteins intermediate values (Kleiber 1975). RQ values can exceed 1.0 when anabolism of fat exceeds catabolism (*i.e.,* de novo lipogenesis (DNL) Benedict 1937; Abreu-Vieira et al. 2015; Levin et al. 2017) or during anaerobic exercise (Whipp 2007; Zagatto et al. 2012). While carbohydrates, fats, or proteins are all burned as fuel, if present in excess, they can also be converted to fat which can be stored for future use (Barboza et al. 2009). DNL is one mechanism for converting non-fat energy into a form that can be stored. The conversion of glucose to fats prior to entry into the citric acid cycle is exothermic and results in an energetic cost, representing a metabolically inefficient use of dietary substrate in cold habitats (Solinas et al. 2015), but a potentially efficient mechanism of heat regulation in hot deserts. RQ was greater than 1.0 during the inactive period in animals on both diets suggesting that *de novo* lipogenesis (DNL) is not limited by fat intake in this species. Here and in previous experiments on *P. eremicus* (Colella et al. 2021b), RQ is only greater than 1.0 during the light phase when animals are inactive, which strongly suggests DNL is occurring. Further, fat storage could be a way for desert mammals to buffer against uncertain water availability, as fat oxidation releases metabolic water (Mellanby 1942; Davidson and Passmore 1963; Schmidt-Nielsen and Adolph 1964). Given this, the conversion of other fuel sources to fats may serve as an important reservoir of metabolic water. Previous studies have shown that DNL can occur under high-carbohydrate and high-fat diets (Strable and Ntambi 2010). This is supported by our results, as both groups had an RQ greater than 1.0 during the light phase and RQ values at or below the FQ during the dark phase, which indicates a shift to the supplied dietary substrate as the main oxidative fuel during cooler, dark periods.

### Electrolytes

Electrolytes provide information on the overall metabolic state, renal function, and other core physiological functions of an organism (Kutscher 1968), including hydration status (Cheuvront et al. 2010) or disease state. Therefore, to further understand the effects of diet on physiological function of small mammals in desert environments, we examined serum electrolytes from male and female cactus mice fed diets with different levels of fat. We established diet-specific differences for serum Na, K, Cr, BUN, Hct, iCa, and osmolality values for *P. eremicus*. Notably, synthetic markers of pathological renal impairment, BUN and Cr, were not significantly different between treatments. Previous studies have shown that the electrolyte-water balance is frequently unaffected when desert rodents are water-deprived for long periods of time (Heimeier et al. 2004; Heimeier and Donald 2006; Boumansour et al. 2021), however, other desert rodent studies have demonstrated evidence of dehydration in the absence of exogenous water without renal impairment (Kordonowy et al. 2017; Boumansour et al. 2021). So, even with the significant water lost through respiration, when free water is available, dietary fat content does not significantly impair the kidney for *P. eremicus*.

Interestingly, other key electrolytes that are sensitive to hydration status were significantly different between the dietary treatment groups (Figure 2). Serum Na, osmolality, and Hct are expected to be elevated in clinical dehydration and were found to be elevated in animals fed the LFD. Though some contrasts were not statistically significant, measurements were consistently higher for both males and females, a strong sign of differences in hydration between treatment groups. These differences are particularly interesting because while animals fed a LFD had higher rates of water loss than those fed the SD, all animals had access to water *ad libitum*. Elevated serum Na, osmolality, and Hct in the context of normal renal function are evidence of dehydration, but dehydration in neuro-intact animals with free access to water is unusual. While dietary fats may be responsible for changes in solute balance (Friedman et al. 2012), typically the relationship is in the opposite direction to what we found (Abdel-Rahman 2010). One possible hypothesis is that there may be an impairment of the physiological mechanisms related to thirst. Thirst is a response to changes in the blood chemistry (*i.e.,* increase in osmolality Gilman 1937; Wolf 1950; Heimeier et al. 2004) or volume (Fitzsimons 1961; Stricker 1966) both of which are monitored by the lamina terminalis circumventricular organs (Bourque 2008; Zimmerman et al. 2017). Activation of these brain regions motivates an organism to find water. Previous studies have found that the expression of two genes, *Rxfp1* and *Pdyn,* in relevant cell types correlates with neural activation under osmotic and hypovolemic thirst in lab mice (Pool et al. 2020). Further studies should examine these genes and others in desert organisms to understand the basis of thirst pathways. Regardless of the process, elevated serum sodium may have important and complex physiological consequences. For example, elevated blood pressure and increased fluid retention as a result of elevated sodium levels, both have obvious implications for survival in desert environments and deserve future study.

## Conclusion

To understand ways in which the desert-adapted cactus mouse, *P. eremicus,* responds to variation in dietary fat content, reproductively mature mice were subjected to one of two diets, a SD or a LFD. Overall, EE and WLR for mice on the SD during the inactive, light phase (a measure comparable to the resting metabolic rate) agreed with earlier studies of *Peromyscus* (Withers 1992; Ramirez et al. 2022). Further, we show that dietary fat is important for water regulation in cactus mice through the analysis of interacting physiological traits: 1) high rates of total water loss during the warmer, drier light phase and 2) a difference in serum electrolyte values indicating dehydration. Our results suggest that a LFD could limit the capacity of desert animals to tolerate limited access to free water as is common in arid environments.

The identification of short-term, physiological responses of *P. eremicus* associated with dietary fat composition demonstrates that adaptations that evolved over long evolutionary timescales will affect survival during the accelerated time scales of climate change. In light of global climate change and increased desertification, investigating the range and mechanisms of plastic responses employed by desert-adapted species could provide insights into physiological responses to increasingly erratic climate.

## Acknowledgments

We would like to thank members of the MacManes Lab for helpful comments and support on early versions of the manuscript; the Animal Resources Office and veterinary care staff at the University of New Hampshire; A. Gerson at the University of Massachusetts Amherst, Z. Cheviron at the University of Montana, B. Joos and J. Klok at Sable Systems International for analytic guidance, technical support, and respirometry training. This work was supported by the National Institute of Health National Institute of General Medical Sciences (R35 GM128843 to M.D.M.).

## Author Contributions

Conceptualization: M.D.M.; Methodology: D.M.B., M.D.M.; Formal analysis: D.M.B., E.L.; Investigation: D.M.B., J.P.C.; Resources: M.D.M.; Writing - original draft: D.M.B.; Writing - review & editing: D.M.B., J.P.C., E.L., M.D.M.; Visualization: D.M.B., E.L.; Supervision: M.D.M.; Project administration: M.D.M.; Funding acquisition: M.D.M.

## Data Availability

Raw ExpeData (SSI) files are available through Zenodo: https://zenodo.org/record/6422231#.YlXSD9PML0o. Macro processing files, raw and processed respirometry data, and cage sampling scheme files are also available on Zenodo. All R scripts used in this project are available through GitHub at: https://github.com/DaniBlumstein/Diet_paper.

## Supplemental

**Table S1:**
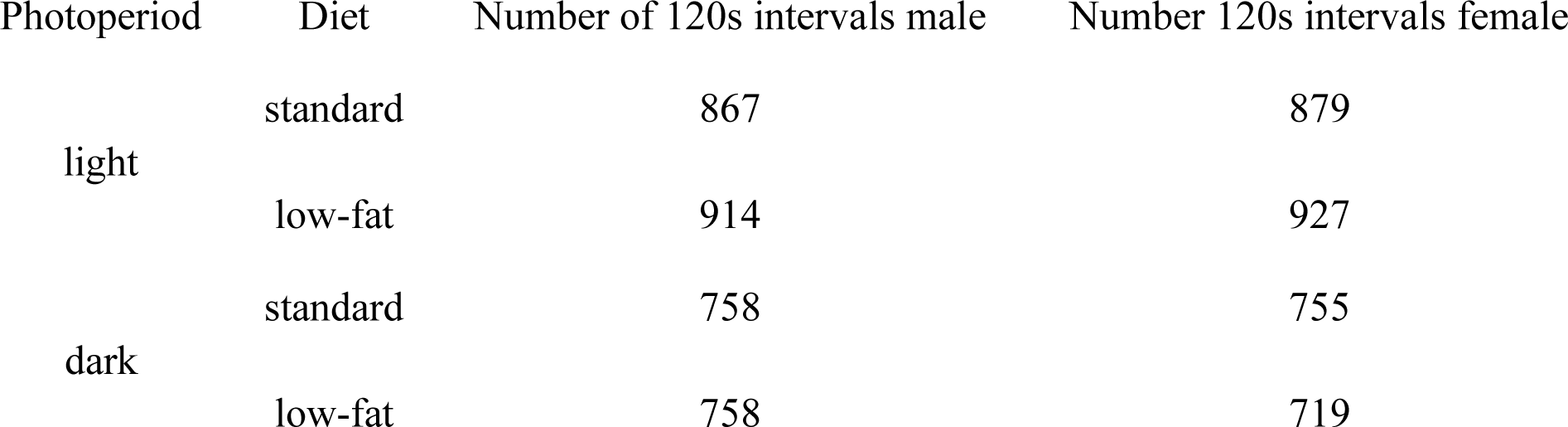
Number of samples collected per time interval.

**Table S2:**
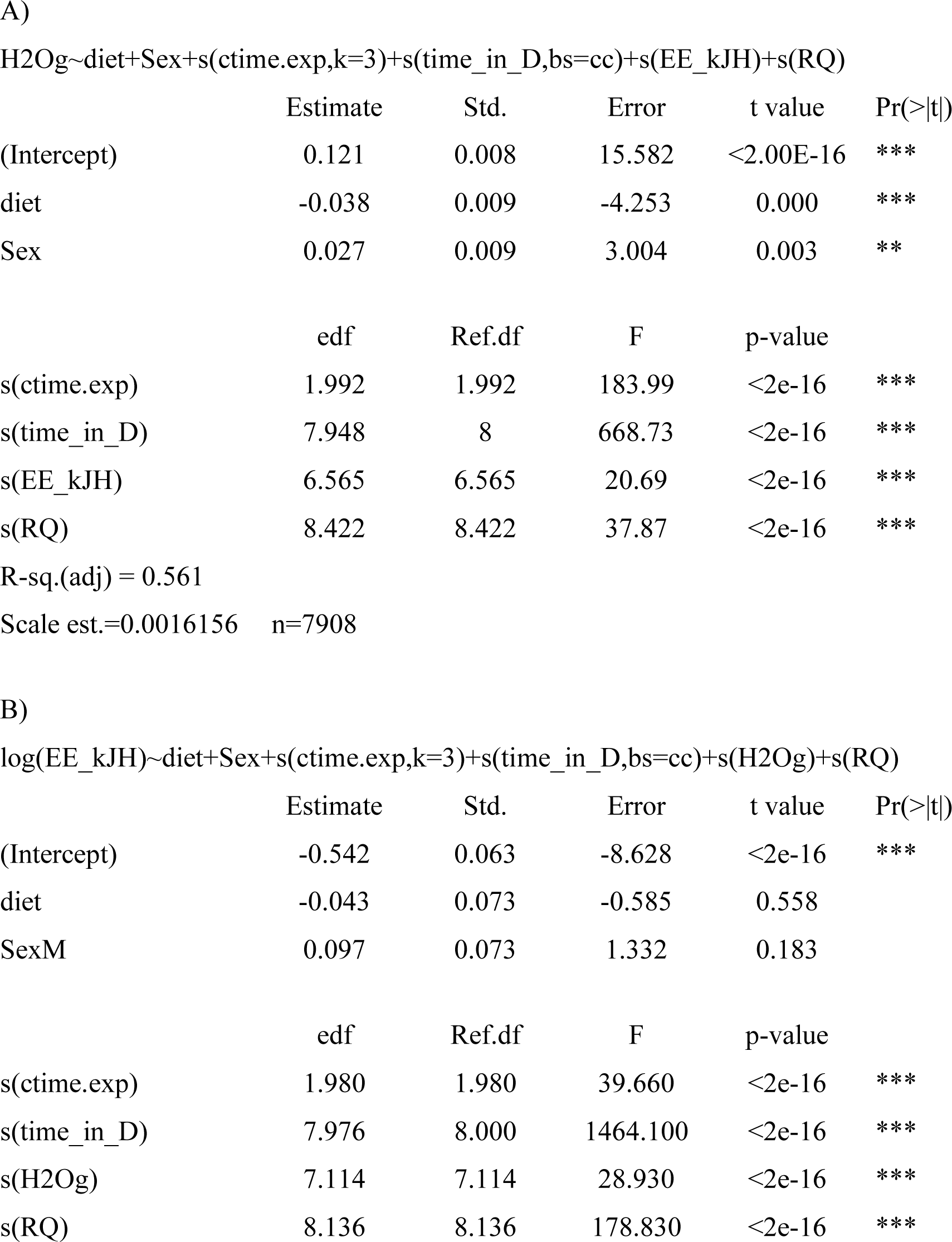
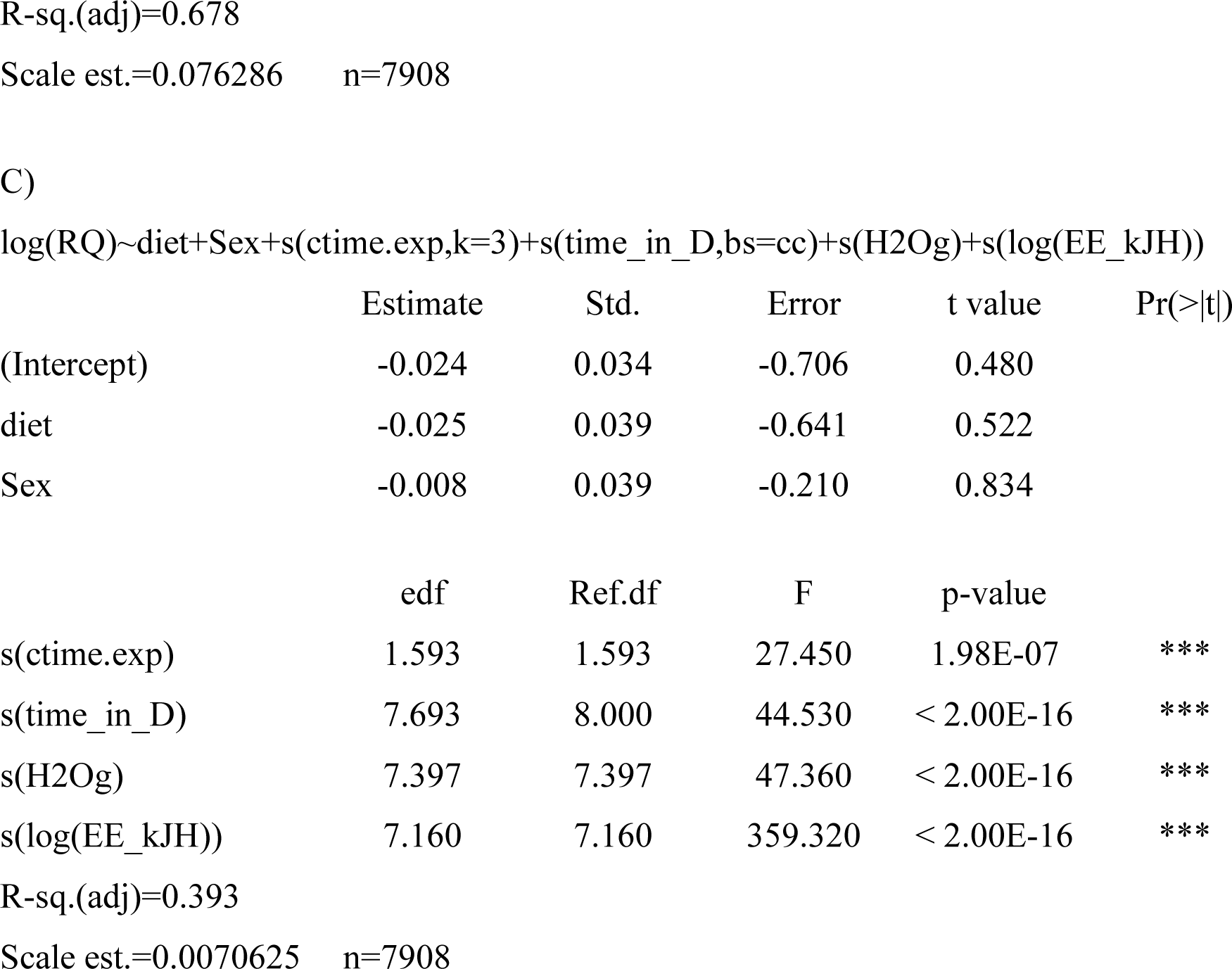
Generalized additive mixed models (GAMM) 24-hour results. Statistical models and results for **A)** water loss rate (WLR, H_2_O g/hr^-1^), **B)** energy expenditure (EE kJ/hr^-1^), **C)** respiratory quotient (RQ).

|  | Estimate | Std. | Error | t value | Pr(>\|t\|) |
| --- | --- | --- | --- | --- | --- |
| (Intercept) | 0.121 | 0.008 | 15.582 | <2.00E-16 | *** |
| diet | -0.038 | 0.009 | -4.253 | 0.000 | *** |
| Sex | 0.027 | 0.009 | 3.004 | 0.003 | ** |
|  | edf | Ref.df | F | p-value |  |
| s(ctime.exp) | 1.992 | 1.992 | 183.99 | <2e-16 | *** |
| s(time_in_D) | 7.948 | 8 | 668.73 | <2e-16 | *** |
| s(EE_kJH) | 6.565 | 6.565 | 20.69 | <2e-16 | *** |
| s(RQ) | 8.422 | 8.422 | 37.87 | <2e-16 | *** |
| R-sq.(adj) = 0.561 |  |  |  |  |  |
| Scale est.=0.0016156 n=7908 |  |  |  |  |  |

B) log(EE\_kJH)~diet+Sex+s(ctime.exp,k=3)+s(time\_in\_D,bs=cc)+s(H2Og)+s(RQ)
|  | Estimate | Std. | Error | t value | Pr(>\|t\|) |
| --- | --- | --- | --- | --- | --- |
| (Intercept) | -0.542 | 0.063 | -8.628 | <2e-16 | *** |
| diet | -0.043 | 0.073 | -0.585 | 0.558 |  |
| SexM | 0.097 | 0.073 | 1.332 | 0.183 |  |
|  | edf | Ref.df | F | p-value |  |
| s(ctime.exp) | 1.980 | 1.980 | 39.660 | <2e-16 | *** |
| s(time_in_D) | 7.976 | 8.000 | 1464.100 | <2e-16 | *** |
| s(H2Og) | 7.114 | 7.114 | 28.930 | <2e-16 | *** |
| s(RQ) | 8.136 | 8.136 | 178.830 | <2e-16 | *** |
| (Intercept) | -0.024 | 0.034 | -0.706 | 0.480 |  |
| diet | -0.025 | 0.039 | -0.641 | 0.522 |  |
| Sex | -0.008 | 0.039 | -0.210 | 0.834 |  |

|  | edf | Ref.df | F | p-value |  |
| --- | --- | --- | --- | --- | --- |
| s(ctime.exp) | 1.593 | 1.593 | 27.450 | 1.98E-07 | *** |
| s(time_in_D) | 7.693 | 8.000 | 44.530 | < 2.00E-16 | *** |
| s(H2Og) | 7.397 | 7.397 | 47.360 | < 2.00E-16 | *** |
| s(log(EE_kJH)) | 7.160 | 7.160 | 359.320 | < 2.00E-16 | *** |

R-sq.(adj)=0.393
Scale est.=0.0070625 n=7908

**Figure S1:**
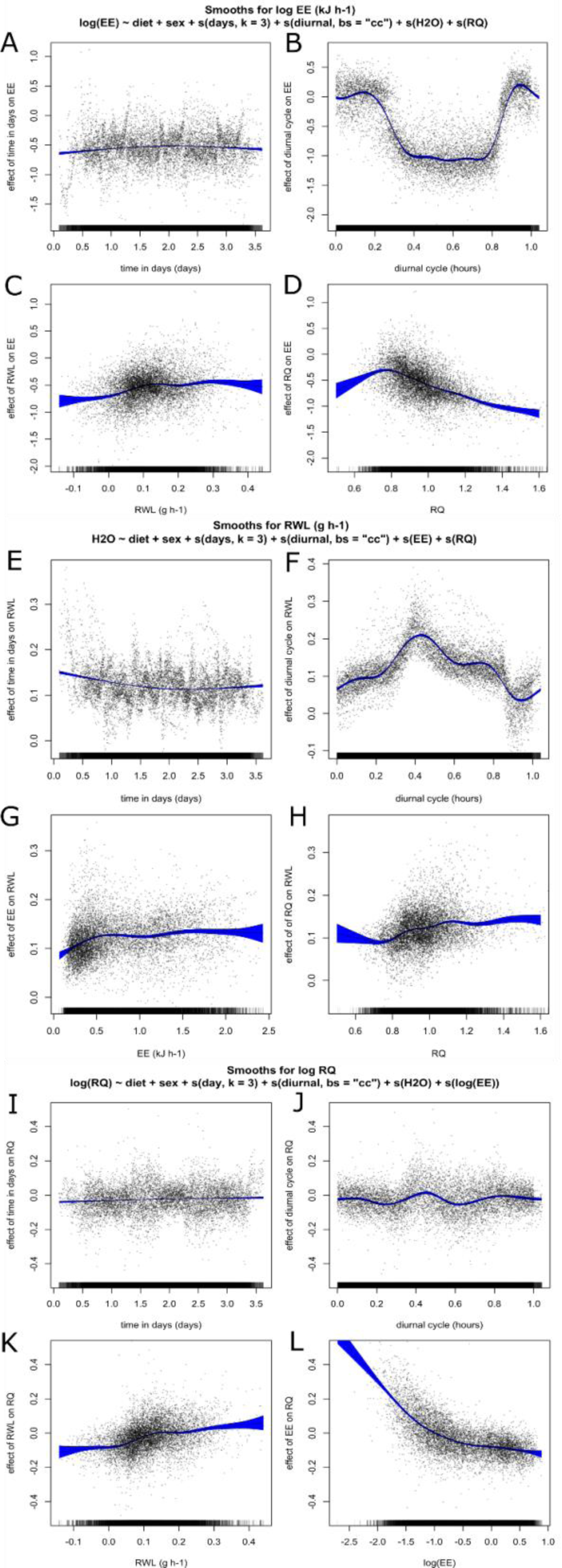
Contributions of model terms to energy expenditure (EE kJ h ^-1^ , A-D), water loss rate (WLR g h ^-1^ , E-H), and respiratory quotient (RQ, I-L) for female and male *Peromyscus eremicus* across two diet groups (standard diet and low-fat diet) in general additive mixed models (GAMM). The smoothing curves for each response variable included two fixed effects; diet (low-fat vs standard) and sex, two random effects; mouse identification number and date of data collection, and four regression terms: time in days, diurnal cycle, and two of the three respirometry response variables (EE, RQ, and/or EWL) as regression terms. For each graph, the y-axis is the effect of the x-axis on the respirometry response variable estimated by a multivariable GAMM. Blue-shaded areas are 95% confidence intervals, black points are residuals.

**Table S3:**
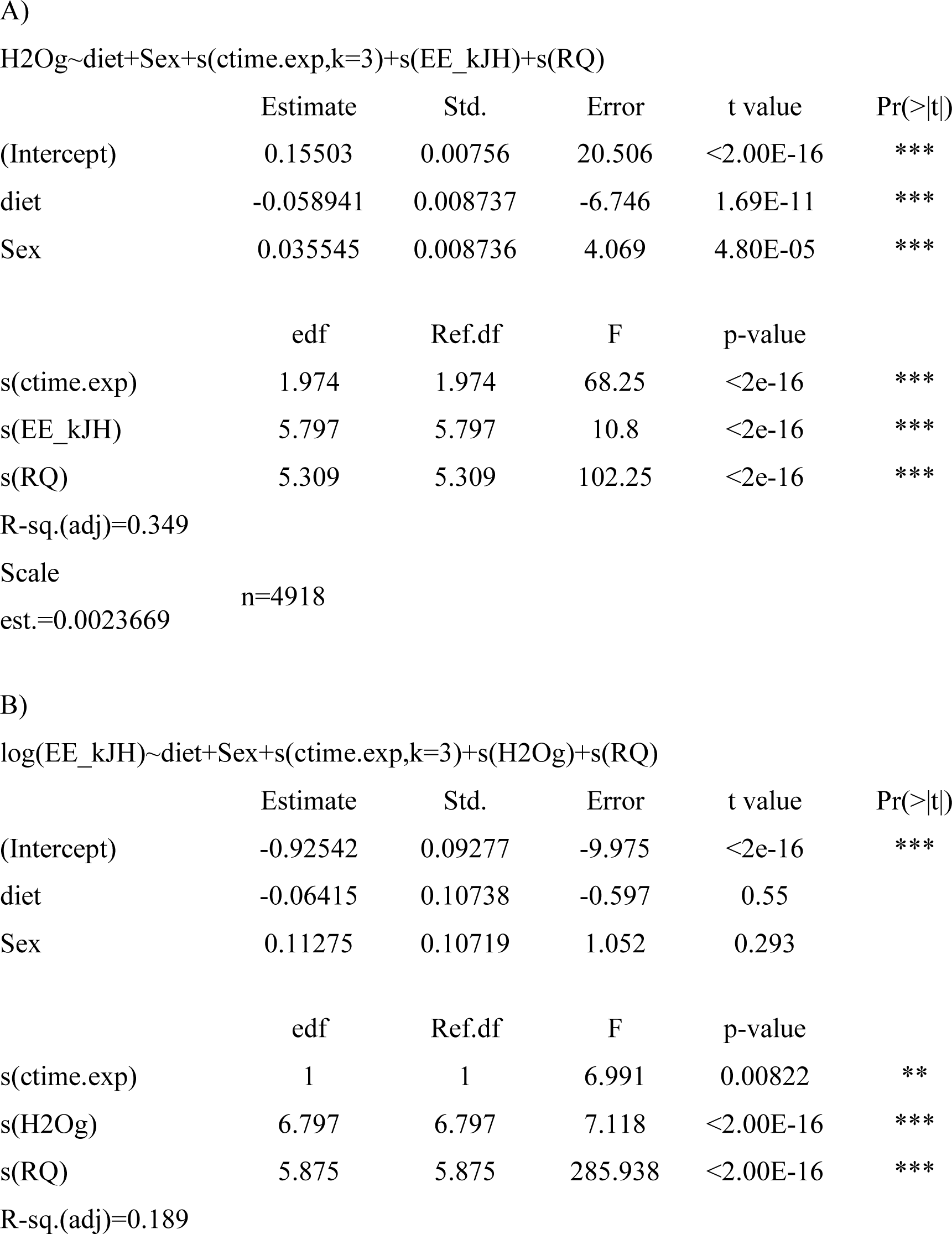
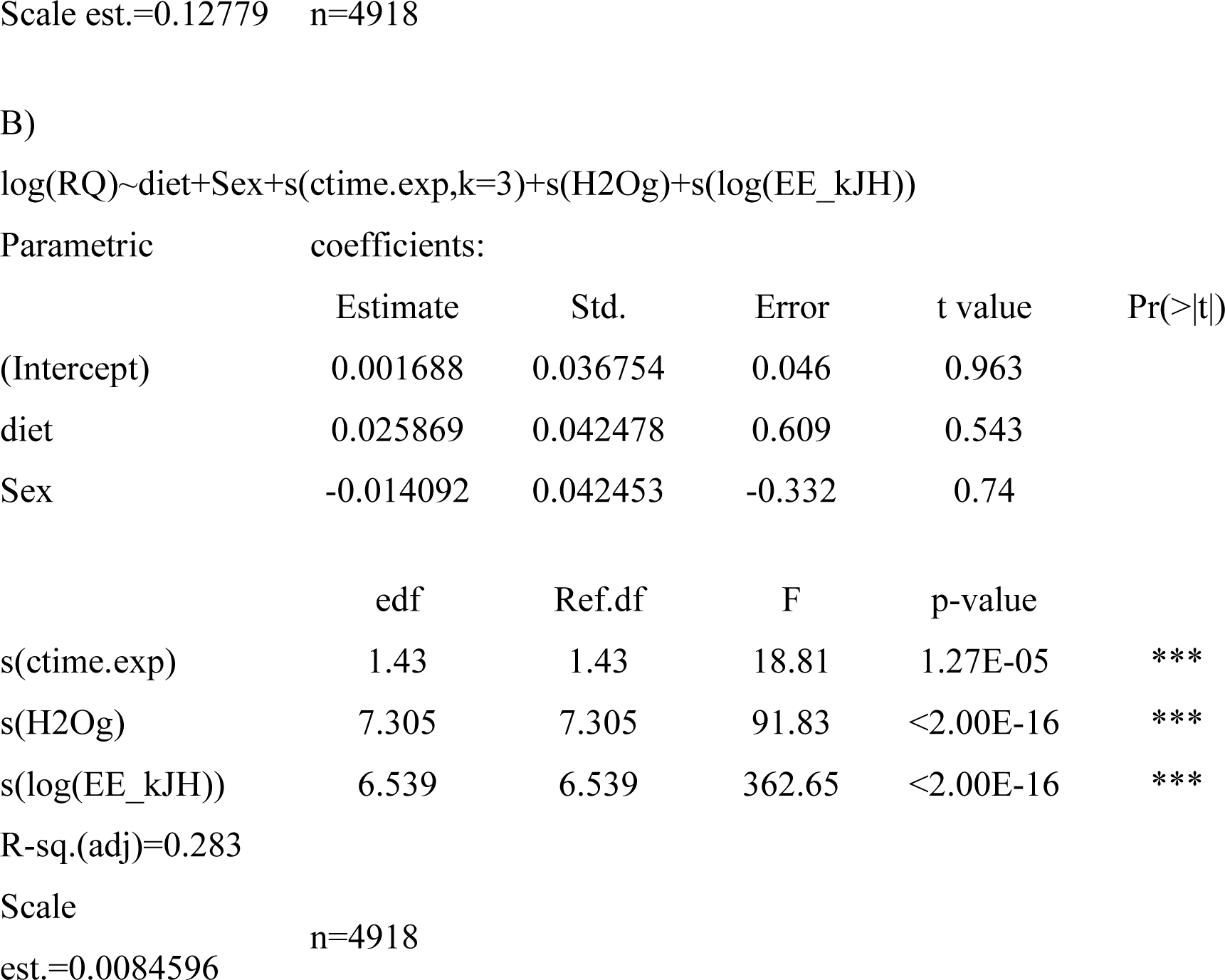
Generalized additive mixed models (GAMM) results for only data collected during the light phase. Statistical models and results for A) water loss rate (WLR, H2O g/hr ^-1^ ), B) energy expenditure (EE kJ/hr ^-1^ ), C) respiratory quotient (RQ).

|  | Estimate | Std. | Error | t value | Pr(>\|t\|) |
| --- | --- | --- | --- | --- | --- |
| (Intercept) | 0.15503 | 0.00756 | 20.506 | <2.00E-16 | *** |
| diet | -0.058941 | 0.008737 | -6.746 | 1.69E-11 | *** |
| Sex | 0.035545 | 0.008736 | 4.069 | 4.80E-05 | *** |

|  | edf | Ref.df | F | p-value |  |
| --- | --- | --- | --- | --- | --- |
| s(ctime.exp) | 1.974 | 1.974 | 68.25 | <2e-16 | *** |
| s(EE_kJH) | 5.797 | 5.797 | 10.8 | <2e-16 | *** |
| s(RQ) | 5.309 | 5.309 | 102.25 | <2e-16 | *** |

|  | Estimate | Std. | Error | t value | Pr(>\|t\|) |
| --- | --- | --- | --- | --- | --- |
| (Intercept) | -0.92542 | 0.09277 | -9.975 | <2e-16 | *** |
| diet | -0.06415 | 0.10738 | -0.597 | 0.55 |  |
| Sex | 0.11275 | 0.10719 | 1.052 | 0.293 |  |

|  | edf | Ref.df | F | p-value |  |
| --- | --- | --- | --- | --- | --- |
| s(ctime.exp) | 1 | 1 | 6.991 | 0.00822 | ** |
| s(H<sub>2</sub>Og) | 6.797 | 6.797 | 7.118 | <2.00E-16 | *** |
| s(RQ) | 5.875 | 5.875 | 285.938 | <2.00E-16 | *** |

| Parametric | coefficients: |  |  |  |  |
| --- | --- | --- | --- | --- | --- |
|  | Estimate | Std. | Error | t value | Pr(>\|t\|) |
| (Intercept) | 0.001688 | 0.036754 | 0.046 | 0.963 |  |
| diet | 0.025869 | 0.042478 | 0.609 | 0.543 |  |
| Sex | -0.014092 | 0.042453 | -0.332 | 0.74 |  |
|  | edf | Ref.df | F | p-value |  |
| s(ctime.exp) | 1.43 | 1.43 | 18.81 | 1.27E-05 | *** |
| s(H2Og) | 7.305 | 7.305 | 91.83 | <2.00E-16 | *** |
| s(log(EE_kJH)) | 6.539 | 6.539 | 362.65 | <2.00E-16 | *** |
| R-sq.(adj)=0.283 |  |  |  |  |  |
| Scale | n=4918 |  |  |  |  |
| est.=0.0084596 |  |  |  |  |  |

**Table S4:** Generalized additive mixed models (GAMM) results for only data collected during the dark phase. Statistical models and results for A) water loss rate (WLR, H2O g/hr ^-1^), B) energy expenditure (EE kJ/hr ^-1^ ), C) respiratory quotient (RQ).

|  | Estimate | Std. | Error | t value | Pr(>\|t\|) |
| --- | --- | --- | --- | --- | --- |
| (Intercept) | 0.053382 | 0.00923 | 5.784 | 8.07E-09 | *** |
| diet | 0.002749 | 0.01071 | 0.257 | 0.797 |  |
| Sex | 0.025882 | 0.010637 | 2.433 | 0.015 | * |

|  | edf | Ref.df | F | p-value |  |
| --- | --- | --- | --- | --- | --- |
| s(ctime.exp) | 1.968 | 1.968 | 19.063 | 4.29E-07 | *** |
| s(EE_kJH) | 4.881 | 4.881 | 8.185 | 6.41E-07 | *** |
| s(RQ) | 7.887 | 7.887 | 47.955 | <2.00E-16 | *** |

|  | Estimate | Std. | Error | t value | Pr(>\|t\|) |
| --- | --- | --- | --- | --- | --- |
| (Intercept) | 0.08701 | 0.07403 | 1.175 | 0.24 |  |
| diet | -0.04856 | 0.0857 | -0.567 | 0.571 |  |
| Sex | 0.11419 | 0.08547 | 1.336 | 0.182 |  |

|  | edf | Ref.df | F | p-value |  |
| --- | --- | --- | --- | --- | --- |
| s(ctime.exp) | 1.97 | 1.97 | 21.35 | <2e-16 | *** |
| s(H<sub>2</sub>Og) | 7.853 | 7.853 | 12.85 | <2e-16 | *** |
| s(RQ) | 7.469 | 7.469 | 34.63 | <2e-16 | *** |

|  | Estimate | Std. | Error | t value |  |
| --- | --- | --- | --- | --- | --- |
| (Intercept) | -0.07254 | 0.02994 | -2.423 | 0.0155 | * |
| diet | -0.08263 | 0.03458 | -2.39 | 0.0169 | * |
| Sex | -0.01154 | 0.03459 | -0.333 | 0.7388 |  |
|  | edf | Ref.df | F | p-value |  |
| s(ctime.exp) | 1.95 | 1.95 | 20.1 | <2e-16 | *** |
| s(H2Og) | 6.454 | 6.454 | 45.5 | <2e-16 | *** |
| s(log(EE_kJH)) | 1 | 1 | 39.76 | <2e-16 | *** |

R-sq.(adj)=0.292
Scale est.=0.0032848
n=2990

**Table S5:** ANCOVA results for total water loss.

|  | Sum | Sq | Df | F value | Pr(>F) |
| --- | --- | --- | --- | --- | --- |
| diet | 130.36 | 1 | 19.5126 | 5.22E-05 | *** |
| sex | 40.4 | 1 | 6.0471 | 0.01736 | * |
| weight | 7.03 | 1 | 1.0529 | 0.30967 |  |
| diet:sex | 1.54 | 1 | 0.2305 | 0.63317 |  |
| Residuals | 340.72 | 51 |  |  |  |
|  | estimate | SE | df | t.ratio | p.value |
| contrast |  |  |  |  |  |
| F Low-fat -<br>M Low-fat | -2.05 | 0.977 | 51 | -2.101 | 0.1667 |
| F Low-fat -<br>F Standard | 2.76 | 1.018 | 51 | 2.716 | 0.0433 |
| F Low-fat -<br>M Standard | 1.39 | 0.977 | 51 | 1.421 | 0.4922 |
| M Low-fat -<br>F Standard | 4.82 | 1.014 | 51 | 4.748 | 0.0001 |
| M Low-fat -<br>M Standard | 3.44 | 0.977 | 51 | 3.522 | 0.0049 |
| F Standard -<br>M Standard | -1.38 | 1.013 | 51 | -1.359 | 0.5307 |

## References

Abdel-Rahman, M. K. 2010. Influence of dietary fat on renal function, lipid profile, sex hormones, and electrolyte balance in rats. European Journal of Lipid Science and Technology 112:1166–1172.

Abreu-Vieira, G., C. Xiao, O. Gavrilova, and M. L. Reitman. 2015. Integration of body temperature into the analysis of energy expenditure in the mouse. Molecular Metabolism 4:461–470.

Acheson, K. J. 1990. Sympathetic Nervous System in the Regulation of Thermogenesis. Pp. 40–46 in Hormones and Nutrition in Obesity and Cachexia (M. J. Müller, E. Danforth, A. G. Burger & U. Siedentopp, eds.). Springer, Berlin, Heidelberg.

Aisenbrey, G. A., W. A. Handelman, P. Arnold, M. Manning, and R. W. Schrier. 1981. Vascular effects of arginine vasopressin during fluid deprivation in the rat. The Journal of Clinical Investigation 67:961–968.

Bachmanov, A. A., D. R. Reed, G. K. Beauchamp, and M. G. Tordoff. 2002. Food Intake, Water Intake, and Drinking Spout Side Preference of 28 Mouse Strains. Behavior genetics 32:435–443.

Barboza, P. S., K. L. Parker, and I. D. Hume (EDS.). 2009. Lipids: Fatty Acids and Adipose Tissue. Pp. 119–131 in Integrative Wildlife Nutrition. Springer, Berlin, Heidelberg.

Beale, P. K., K. J. Marsh, W. J. Foley, and B. D. Moore. 2018. A hot lunch for herbivores: physiological effects of elevated temperatures on mammalian feeding ecology. Biological Reviews 93:674–692.

Bedford, N. L., and H. E. Hoekstra. 2015. Peromyscus mice as a model for studying natural variation. eLife 4:e06813.

Benedict, F. G. 1937. Lipogenesis in the animal body, with special reference to the physiology of the goose. Carnegie institution of Washington.

Boogert, N. J., J. R. Madden, J. Morand-Ferron, and A. Thornton. 2018. Measuring and understanding individual differences in cognition. Philosophical Transactions of the Royal Society B: Biological Sciences 373:20170280.

Boumansour, L., et al. 2021. Vasopressin and oxytocin expression in hypothalamic supraoptic nucleus and plasma electrolytes changes in water-deprived male Meriones libycus. Animal Cells and Systems 0:1–10.

Bourque, C. W. 2008. Central mechanisms of osmosensation and systemic osmoregulation. Nature Reviews Neuroscience 9:519–531.

Bradley, W. G., AND R. A. Mauer. 1973. Rodents of a Creosote Bush Community in Southern Nevada. The Southwestern Naturalist 17:333–344.

Calder, W. A., AND E. J. Braun. 1983. Scaling of osmotic regulation in mammals and birds. The American Journal of Physiology 244:R601–606.

Castle, S. T., ET AL. 2020. Diet composition analysis provides new management insights for a highly specialized endangered small mammal. PLOS ONE 15:e0240136.

Cheuvront, S. N., R. W. Kenefick, S. J. Montain, and M. N. Sawka. 2010. Mechanisms of aerobic performance impairment with heat stress and dehydration. Journal of Applied Physiology 109:1989–1995.

Colella, J. P., et al. 2021a. Limited Evidence for Parallel Evolution Among Desert-Adapted Peromyscus Deer Mice. Journal of Heredity 112:286–302.

Colella, J. P., D. M. Blumstein, and M. D. Macmanes. 2021b. Disentangling environmental drivers of circadian metabolism in desert-adapted mice. Journal of Experimental Biology 224.

Crossland, J. P., M. J. Dewey, S. C. Barlow, P. B. Vrana, M. R. Felder, and G. J. Szalai. 2014. Caring for Peromyscus spp. in research environments. Lab Animal 43:162–166.

Davidson, S., and R. Passmore. 1963. Human nutrition and dietetics. Human nutrition and dietetics.

Fitzsimons, J. T. 1961. Drinking by rats depleted of body fluid without increase in osmotic pressure. The Journal of Physiology 159:297.

Fox, J., and S. Weisberg. 2019. An {R} Companion to Applied Regression. Third. Sage. Frank, C. L. 1988. Diet Selection by a Heteromyid Rodent: Role of Net Metabolic Water Production. Ecology 69:1943–1951.

Freeman, J., P. Veggiotti, G. Lanzi, A. Tagliabue, and E. Perucca. 2006. The ketogenic diet: from molecular mechanisms to clinical effects. Epilepsy Res 68:145–80.

Friedman, A. N., et al. 2012. Comparative effects of low-carbohydrate high-protein versus low- fat diets on the kidney. Clinical journal of the American Society of Nephrology: CJASN 7:1103–1111.

Garland, T., JR, ET AL. 2011. The biological control of voluntary exercise, spontaneous physical activity and daily energy expenditure in relation to obesity: human and rodent perspectives. Journal of Experimental Biology 214:206–229.

Gilman, A. 1937. The relation between blood osmotic pressure, fluid distribution and voluntary water intake. American Journal of Physiology-Legacy Content 120:323–328.

Greenleaf, J. E. 1992. Problem: thirst, drinking behavior, and involuntary dehydration. Medicine & Science in Sports & Exercise 24:645.

Hart, J. S. 1971. Rodents. Pp. 1–149 in Mammals. Elsevier.

Heimeier, R. A., R. C. Bartolo, and J. A. Donald. 2004. The effect of water deprivation on signalling molecules that utilise cGMP in the spinifex hopping mouse Notomys alexis. Australian Mammalogy 26:191–198.

Heimeier, R. A., and J. A. Donald. 2006. The response of the natriuretic peptide system to water deprivation in the desert rodent, Notomys alexis. Comparative Biochemistry and Physiology Part A: Molecular & Integrative Physiology 143:193–201.

Hope, A., and R. Parmenter. 2007. Food Habits of Rodents Inhabiting Arid and Semi-arid Ecosystems of Central New Mexico. Special Publications.

Jiang, P., et al. 2012. Major taste loss in carnivorous mammals. Proceedings of the National Academy of Sciences 109:4956–4961.

Kaiyala, K. J., et al. 2012. Acutely Decreased Thermoregulatory Energy Expenditure or Decreased Activity Energy Expenditure Both Acutely Reduce Food Intake in Mice. PLOS ONE 7:e41473.

Kishida, T., J. Thewissen, T. Hayakawa, H. Imai, AND K. Agata. 2015. Aquatic adaptation and the evolution of smell and taste in whales. Zoological Letters 1:9.

Kleiber, M. 1975. The fire of life. An introduction to animal energetics.

Kordonowy, L. et al. 2017. Physiological and biochemical changes associated with acute experimental dehydration in the desert adapted mouse, *Peromyscus eremicus*. Physiological Reports 5:e13218.

Kutscher, C. 1968. Plasma volume change during water-deprivation in gerbils, hamsters, guinea pigs and rats. Comparative Biochemistry and Physiology 25:929–936.

Leary, S. L. et al. 2013. AVMA guidelines for the euthanasia of animals: 2013 edition. American Veterinary Medical Association Schaumburg, IL.

Leib, D. E., C. A. Zimmerman, and Z. A. Knight. 2016. Thirst. Current Biology 26:R1260–R1265.

Levin, E., G. Lopez-Martinez, B. Fane, and G. Davidowitz. 2017. Hawkmoths use nectar sugar to reduce oxidative damage from flight. Science 355:733–735.

Lighton, J. R. B. 2018. Measuring Metabolic Rates: A Manual for Scientists. 2nd edition. Oxford University Press.

Lin, X., and D. Zhang. 1999. Inference in Generalized Additive Mixed Models by Using Smoothing Splines. Journal of the Royal Statistical Society. Series B (Statistical Methodology) 61:381–400.

Macmanes, M. D. 2017. Severe acute dehydration in a desert rodent elicits a transcriptional response that effectively prevents kidney injury. Renal Physiol:11.

Macmillen, R. E. 1965. Aestivation in the cactus mouse, Peromyscus eremicus. Comparative Biochemistry and Physiology 16:227–248.

Macmillen, R. E. 1972. Water economy of nocturnal desert rodents.

Mcarthur, E. D. 1994. Nutritive Quality and Mineral Content of Potential Desert Tortoise Food Plants. Intermountain Research Station.

Mellanby, K. 1942. Metabolic Water and Desiccation. Nature 150:21–21.

Meserve, P. L. 1976. Habitat and Resource Utilization by Rodents of a California Coastal Sage Scrub Community. Journal of Animal Ecology 45:647–666.

Murie, M. 1961. Metabolic Characteristics of Mountain, Desert and Coastal Populations of Perormyscus. Ecology 42:723–740.

Noble, J. D., S. L. Collins, A. J. Hallmark, K. Maldonado, B. O. Wolf, AND S. D. Newsome. 2019. Foraging strategies of individual silky pocket mice over a boom–bust cycle in a stochastic dryland ecosystem. Oecologia 190:569–578.

Orr, T. J., S. D. Newsome, and B. O. Wolf. 2015. Cacti supply limited nutrients to a desert rodent community. Oecologia 178:1045–1062.

Pardi, M. I., R. C. Terry, E. A. Rickart, and R. J. Rowe. 2020. Testing climate tracking of montane rodent distributions over the past century within the Great Basin ecoregion. Global Ecology and Conservation 24:e01238.

Pergams, O. R. W., and J. J. Lawler. 2009. Recent and Widespread Rapid Morphological Change in Rodents. PLOS ONE 4:e6452.

Pool, A.-H., et al. 2020. The cellular basis of distinct thirst modalities. Nature 588:112–117.

Porter, W. P., and D. M. Gates. 1969. Thermodynamic equilibria of animals with environment. Ecological monographs 39:227–244.

Pyke, G. H. 1984. Optimal Foraging Theory: A Critical Review. Annual Review of Ecology and Systematics 15:523–575.

Pyke, G. H., H. R. Pulliam, and E. L. Charnov. 1977. Optimal Foraging: A Selective Review of Theory and Tests. The Quarterly Review of Biology 52:137–154.

R Core Team. 2020. R: A language and environment for statistical computing. R Foundation for Statistical Computing, Vienna, Austria. URL https://www.R-project.org/.

Ramirez, R. W., E. A. Riddell, S. R. Beissinger, and B. O. Wolf. 2022. Keeping your cool: thermoregulatory performance and plasticity in desert cricetid rodents. Journal of Experimental Biology 225:jeb243131.

Rasouli, M. 2016. Basic concepts and practical equations on osmolality: Biochemical approach. Clinical Biochemistry 49:936–941.

Reichman, O. J. 1975. Relation of Desert Rodent Diets to Available Resources. Journal of Mammalogy 56:731–751.

Russell V. Lenth. 2021. emmeans: Estimated Marginal Means, aka Least-Squares Means. R package version 1.6.0. https://CRAN.R-project.org/package=emmeans.

Sánchez-Peña, M. J., F. Márquez-Sandoval, A. C. Ramírez-Anguiano, S. F. Velasco- Ramírez, G. Macedo-Ojeda, and L. J. González-Ortiz. 2017. Calculating the metabolizable energy of macronutrients: a critical review of Atwater’s results. Nutrition Reviews 75:37–48.

Schmidt-Nielsen, B., AND K. SCHMIDT-NIELSEN. 1951. A complete account of the water metabolism in kangaroo rats and an experimental verification. Journal of Cellular and Comparative Physiology 38:165–181.

Schmidt-Nielsen, K. 1975. Desert Rodents: Physiological Problems of Desert Life. Pp. 379– 388 in Rodents in Desert Environments (I. Prakash & P. K. Ghosh, eds.). Springer Netherlands, Dordrecht.

Schmidt-Nielsen, K., and E. F. Adolph. 1964. Desert Animals: Physiological Problems of Heat and Water. Physiological Zoology 37:338–339.

Secor, S. M. 2009. Specific dynamic action: a review of the postprandial metabolic response. Journal of Comparative Physiology B 179:1–56.

Sikes, R. S., and the Animal Care and Use Committee of the American Society of MAMMALOGISTS. 2016. 2016 Guidelines of the American Society of Mammalogists for the use of wild mammals in research and education. Journal of Mammalogy 97:663–688.

Solinas, G., J. Borén, and A. G. Dulloo. 2015. De novo lipogenesis in metabolic homeostasis: More friend than foe? Molecular Metabolism 4:367–377.

Speakman, J. 1997. Factors influencing the daily energy expenditure of small mammals. Proceedings of the Nutrition Society 56:1119–1136.

Strable, M. S., and J. M. Ntambi. 2010. Genetic control of de novo lipogenesis: role in diet- induced obesity. Critical reviews in biochemistry and molecular biology 45:199–214.

Stricker, E. M. 1966. Extracellular fluid volume and thirst. American Journal of Physiology- Legacy Content 211:232–238.

Thornton, S. N. 2010. Thirst and hydration: Physiology and consequences of dysfunction. Physiology & Behavior 100:15–21.

Tigano, A., J. P. Colella, and M. D. Macmanes. 2020. Comparative and population genomics approaches reveal the basis of adaptation to deserts in a small rodent. Molecular Ecology 29:1300–1314.

Veal, R., and W. Caire. 1979. Peromyscus eremicus. Mammalian Species:1–6.

Westerterp, K. R. 2004. Diet induced thermogenesis. Nutrition & metabolism 1:1–5.

Wheless, J. W. 2001. The ketogenic diet: an effective medical therapy with side effects. Journal of Child Neurology 16:633–635.

Whipp, B. J. 2007. Physiological mechanisms dissociating pulmonary CO2 and O2 exchange dynamics during exercise in humans. Experimental Physiology 92:347–355.

Whitfield, M. C., B. Smit, A. E. Mckechnie, and B. O. Wolf. 2015. Avian thermoregulation in the heat: scaling of heat tolerance and evaporative cooling capacity in three southern African arid-zone passerines. Journal of Experimental Biology 218:1705–1714.

Withers, P. C. 1982. Effect of diet and assimilation efficiency on water balance for two desert rodents. Journal of Arid Environments 5:375–384.

Withers, P. C. 1992. Comparative animal physiology. Saunders College Pub. Philadelphia. Wolf, A. V. 1950. Osmometric analysis of thirst in man and dog. American Journal of Physiology-Legacy Content 161:75–86.

Wolf, B. O., and C. M. Del Rio. 2003. How important are columnar cacti as sources of water and nutrients for desert consumers? A review. Isotopes in Environmental and Health Studies 39:53–67.

Wood, S. 2017. Generalized Additive Models: An Introduction with R, Second Edition. Zagatto, A. M., W. E. Miyagi, R. L. Sakugawa, E. I. Kaminagakura, and M. Papoti. 2012. Aerobic Endurance Measurement by Respiratory Exchange Ratio during a Cycle Ergometer Graded Exercise Test. Journal of Exercise Physiology Online 15:49–56.

Zimmerman, C. A., D. E. Leib, and Z. A. Knight. 2017. Neural circuits underlying thirst and fluid homeostasis. Nature Reviews Neuroscience 18:459–469.

