## Supplementary figures and images for "High total water loss driven by low-fat diet in desert-adapted mice"

### Supplemental figure 1

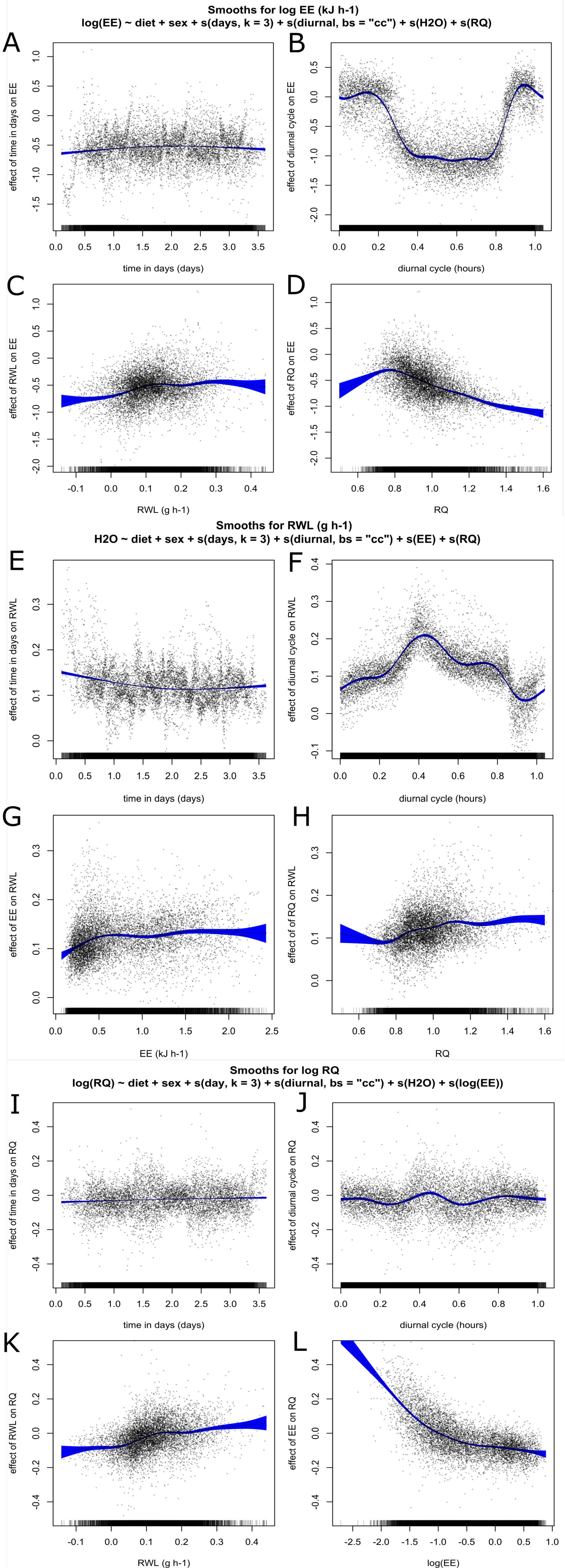
